# Influenza viral particles harboring the SARS-CoV-2 spike RBD as a combination respiratory disease vaccine

**DOI:** 10.1101/2021.04.30.441968

**Authors:** Ryan R. Chaparian, Alfred T. Harding, Kristina Riebe, Amelia Karlsson, Gregory D. Sempowski, Nicholas S. Heaton, Brook E. Heaton

## Abstract

Vaccines targeting SARS-CoV-2 have gained emergency FDA approval, however the breadth against emerging variants and the longevity of protection remains unknown. Post-immunization boosting may be required, perhaps on an annual basis if the virus becomes an endemic pathogen. Seasonal influenza virus vaccines are already developed every year, an undertaking made possible by a robust global vaccine production and distribution infrastructure. To create a seasonal combination vaccine targeting influenza viruses and SARS-CoV-2 that is also amenable to frequent reformulation, we have developed a recombinant influenza A virus (IAV) genetic platform that “reprograms” the virus to package an immunogenic domain of the SARS-CoV-2 spike (S) protein onto IAV particles. Vaccination with this combination vaccine elicits neutralizing antibodies and provides protection from lethal challenge with both pathogens. This technology may allow for leveraging of established influenza vaccine infrastructure to generate a cost-effective and scalable seasonal vaccine solution for both influenza and coronaviruses.

## Introduction

Every year, sufficient influenza virus vaccines to vaccinate the global population are produced, distributed, and administered^1^. These vaccines are widely accepted to be safe and efficacious, however they must be constantly reformatted due to viral antigenic drift^2, 3, 4^. Additionally, a robust reverse genetics system for influenza viruses has been developed, allowing for precise manipulation of the influenza viral genome^5^. Reverse genetics has allowed for the introduction of non-influenza proteins and immune epitopes into influenza viral strains^6, 7, 8^. Thus, at least in theory, leveraging existing influenza virus vaccine production infrastructure to produce recombinant viral strains that express immunogenic antigens from other pathogens of concern may be a practical approach to generating cost-effective, easily implemented combination vaccines or boosters.

SARS-CoV-2 is a respiratory RNA virus that causes COVID-19 disease that is similar in many respects to influenza-virus induced disease^9^. While a number of vaccines designed to vaccinate immunologically naïve people and provide protection from COVID-19 are either in use under emergency authorization from the FDA or in advanced stages of development, these vaccines are for the most part: expensive, associated with higher-than-standard side effects, and difficult to produce/distribute^10, 11, 12, 13^. Further complicating vaccination efforts is the emergence of variant or mutant strains of SARS-CoV-2 that have been associated with reduced protective vaccine efficacy^14^. Additionally, protective immunity against human coronaviruses in general, is thought to be relatively short lived^15, 16, 17, 18^. Thus, it is likely that a cost-effective, scalable, and safe vaccine to periodically boost immunity against SARS-CoV-2 will be needed.

In order to develop a platform-based solution to regularly boost immunity against SARS-CoV-2 and associated variants, we have developed and tested a combination influenza virus-based vaccine that can incorporate both IAV and SARS-CoV-2 antigens. By generating a replication competent IAV that encodes, stably expresses, and packages a key immunogenic domain of the SARS-CoV-2 S protein, we have generated a combination vaccine that can be manufactured in the exact same way most influenza virus vaccines are already produced. To limit cost and the human vaccine burden, a seasonal IAV/SARS-CoV-2 combination vaccine could be used to replace the standard seasonal IAV vaccine and provide protection against novel variants of IAV and SARS-CoV-2. Such an approach would not require additional vaccine manufacturing or distribution facilities, and the resulting effect would be the simultaneous boosting of immunity to two respiratory pathogens.

## Results

### Generation of an H1N1 IAV encoding the SARS-CoV-2 RBD

Our initial goal was to develop a strain of influenza A virus (IAV) that would encode a gene for the receptor binding domain (RBD) of the major antigen of SARS-CoV-2, the spike (S) glycoprotein^19, 20^. We wanted this gene to be expressed at high levels and for the translated SARS-CoV-2 antigen to be packaged onto the IAV particle such that the resulting vaccine could be administered as either live attenuated or inactivated vaccines, formulations both currently approved for use in humans^21^. We have previously published viral genetic modification strategies that allows for the incorporation of a foreign open reading frame into the viral genome^22^. Using the classical A/Puerto Rico/8/1934 H1N1 IAV strain as a proof-of-concept platform, we selected one viral locus, 5’ to the viral hemagglutinin (HA) ORF, for insertion of the SARS-CoV-2 RBD (**Figure 1A, top**) as it is associated with high expression of a foreign gene^22^. In order to facilitate incorporation of the SARS-CoV-2 RBD onto nascent viral particles, we fused the ORF to a specific N-terminal sequence from the IAV neuraminidase (NA) protein which contains the transmembrane domain and cytoplasmic tail of that protein. In between the two proteins is a picornaviral 2A motif which ensures that the proteins are co-translationally separated and can be independently packaged onto the viral particle (**Figure 1A, bottom**).

**Figure 1.**
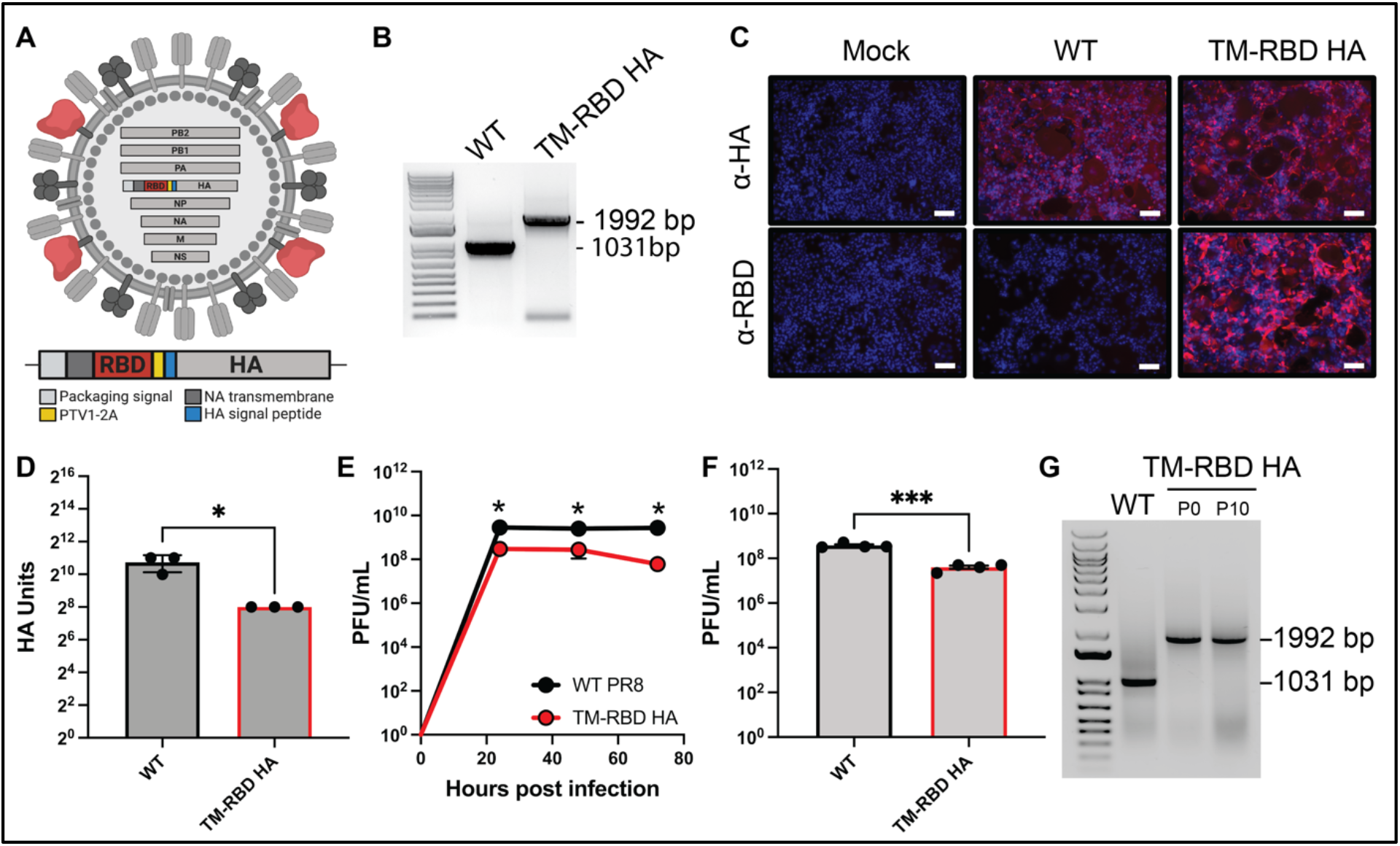
Generation of an IAV strain that stably incorporates the SARS-CoV-2 spike RBD domain into the viral particle. (A) Diagram showing genetic modulation of the HA segment to enable insertion of a foreign ORF. The SARS-CoV-2 RBD was fused to the NA transmembrane domain and a PTV1-2A site was introduced to allow for co-translation of the RBD and HA. (B) Segment 4 RT-PCRs with WT and TM-RBD HA viruses. (C) Immunofluorescence microscopy images of non-permeablized cells infected with WT or TM-RBD virus and stained for HA (top) or SARS-CoV-2 RBD (bottom). Scale bar indicates 100 μm. (D) HA assays of WT/TM-RBD HA viruses after growth in embryonated chicken eggs for 72 hours. (E) Plaque assays of WT/TM-RBD HA viruses grown in embryonated chicken eggs at 24, 48, and 72 hours. (F) Plaque assays WT/TM-RBD HA viruses of after growth in embryonated chicken eggs for 72 hours. (G) Segment 4 RT-PCRs with WT and TM-RBD HA viruses after 10 passages on MDCK cells. P0 indicates the stock of virus used for the experiment. Statistical analyses were performed using unpaired t-tests. For all panels, P-values denoted with asterisks correspond to the following values: * < 0.05, *** < 0.001, and ns = not significant.

We rescued the virus and verified appropriate insertion of the SARS-CoV-2 RBD ORF into the HA-encoding IAV segment via RT-PCR (**Figure 1B**). To determine if the RBD protein was being expressed and localized appropriately to the plasma membrane, we infected cells and applied antibodies against the IAV HA protein and the SARS-CoV-2 RBD to non-permeabilized cells. While infection with both non-modified wild-type (WT) IAV as well as the recombinant vaccine strain (TM-RBD HA) led to expression of HA to the cell surface, membrane-anchored SARS-CoV-2 RBD was only detectable after infection with the recombinant virus (**Figure 1C**). Further, the SARS-CoV-2 RBD expressing virus was able to grow to high titers and with similar kinetics to WT virus (**Figure 1D-F**). Additionally, we wanted to ensure that the SARS-CoV-2 ORF would remain stable in the virus genome. After 10 serial passages, the SARS-CoV-2 RBD ORF remained detectable with no appreciable change in RT-PCR product (**Figure 1G**). Thus, we have demonstrated that we can engineer influenza viruses to stably express a membrane-anchored foreign protein (the SARS-CoV-2 RBD) that grows to levels similar to those of unmodified influenza viruses.

### The SARS-CoV-2 RBD is packaged into IAV particles without disruption of native viral proteins

Next, we wanted to ensure that the membrane-anchored SARS-CoV-2 RBD was in fact incorporated into the viral particle. We therefore purified WT IAV and the recombinant TM-RBD HA viruses and analyzed their protein composition via western blot. We first normalized loading based on the levels of the IAV matrix (M1) protein in the two samples. The M1 protein forms the viral capsid and is not expected to be altered by changes to glycoprotein composition^23^. As expected, we observed that SARS-CoV-2 RBD was detectable in the recombinant RBD virus, but not in the WT virus (**Figure 2A**). In order to determine the influence of the incorporation of another protein in the viral envelope on the native influenza viral surface proteins, we probed for expression of M2 as well as the HA and NA proteins. While the levels of the M2 protein were unchanged, there was a slight reduction in the amount of HA and an increase in the amount of NA packaged into nascent virions compared to WT virus (**Figure 2A**). Thus, this genetic approach can facilitate packaging of a foreign protein onto a viral particle without dramatic effects on native viral protein incorporation.

**Figure 2.**
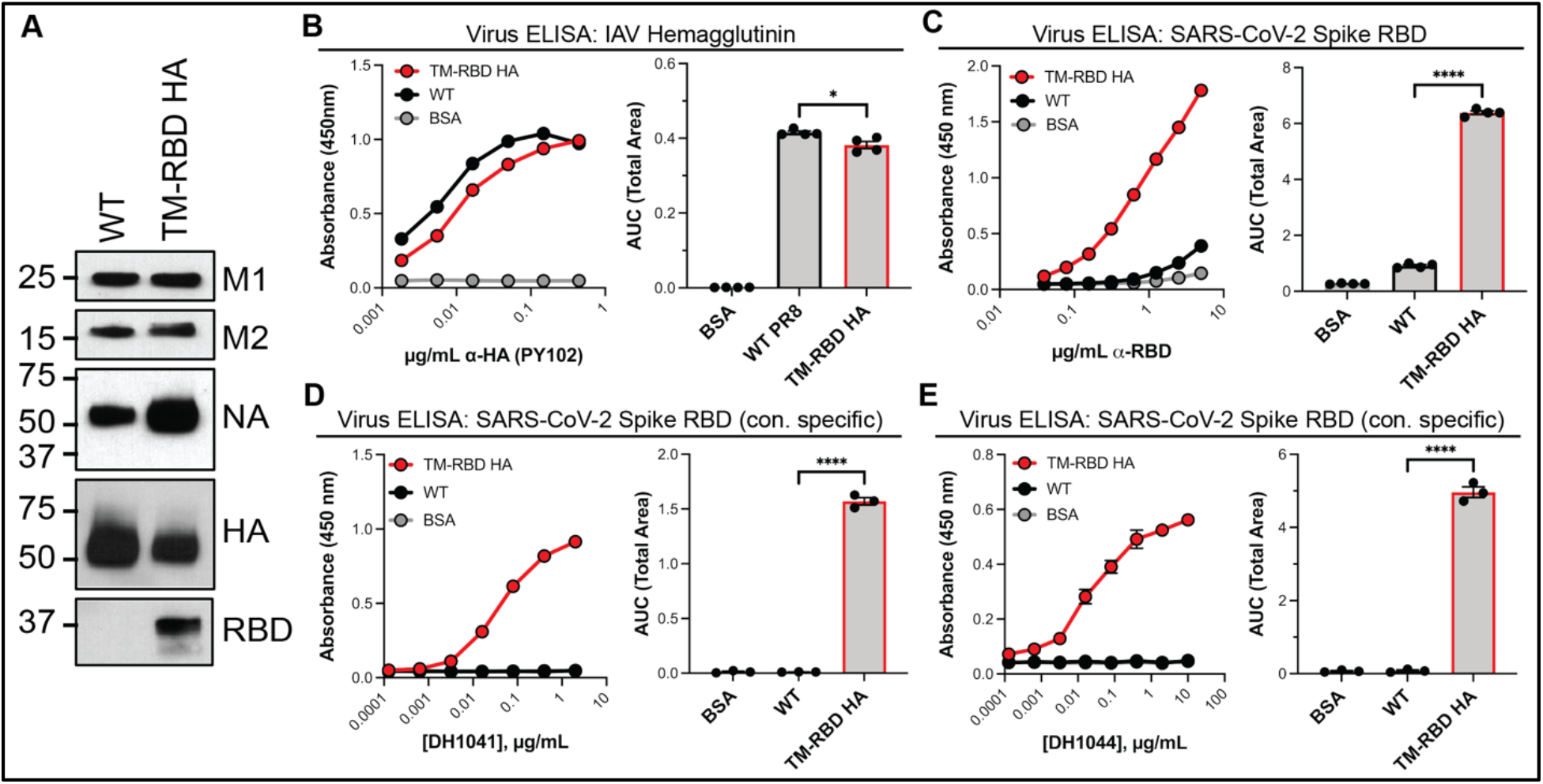
The TM-RBD-HA virus incorporates appropriately folded RBD without disrupting incorporation of the IAV viral envelope proteins. (A) Western blot analysis of WT and TM-RBD HA viruses. Samples were normalized via M1 protein signal using pixel densitometry. (B) (left) ELISAs using the PY102 anti-HA antibody against whole virus particles and (right) area under the curve analysis. (C) (left) ELISAs using a SARS-CoV-2 anti-RBD antibody (binds a non-structural epitope) against whole virus particles and (right) area under the curve analysis. (D) ELISAs using a SARS-CoV-2 neutralizing antibody (DH1041, binds a structural epitope on the RBD), against whole virus particles and (right) area under the curve analysis. (E) Same analysis as in (D) using a different conformation-specific SARS-CoV-2 neutralizing antibody, DH1044. Statistical analyses were performed using unpaired t-tests. For all panels, P-values denoted with asterisks correspond to the following values: * < 0.05, **** < 0.0001, and ns = not significant.

We also wanted to assess the protein packaging under non-denaturing conditions by performing enzyme-linked immunosorbent assays (ELISAs) with fully intact virions. Confirming our western blot results, we observed a statistically significant but minor reduction of HA in the recombinant TM-RBD HA virus compared to unmodified WT virus (**Figure 2B**). We further probed for the SARS-CoV-2 RBD using antibodies that recognize either linear or structural epitopes. A non-conformation specific antibody raised against the RBD yielded strong signal only against the RBD virus (**Figure 2C**). To test if the RBD-TM fusion protein was folding correctly, we utilized DH1041 and DH1044 which are conformation specific human monoclonal antibodies that bind to the SARS-CoV-2 RBD and also display neutralizing abilities^24^. Both of these antibodies specifically bound the TM-RBD HA virus and not the parental WT IAV (**Figure 2D-E**). Taken together, these results indicate that the recombinant TM-RBD HA virus packages properly folded SARS-CoV-2 RBD protein while maintaining the packaging of the other IAV envelope proteins HA, NA, and M2.

### The H1N1/SARS-CoV-2 combination vaccine elicits IAV and SARS-CoV-2 neutralizing antibodies

Our next step was to define if the recombinant IAV/SARS-CoV-2 virus elicited the desired immune responses after vaccination. We therefore took naïve mice and primed them with a live vaccination of either WT or the combination IAV/SARS-CoV-2 viral vaccines. Three weeks later, the animals were boosted with an IM injection of the inactivated vaccine preps or BSA as a negative control. Two weeks after the boost, peripheral blood was collected and serum reactivity against the IAV HA protein or the SARS-CoV-2 RBD protein, as well as the neutralizing viral titers against IAV and SARS-CoV-2, were determined (**Figure 3A**).

**Figure 3.**
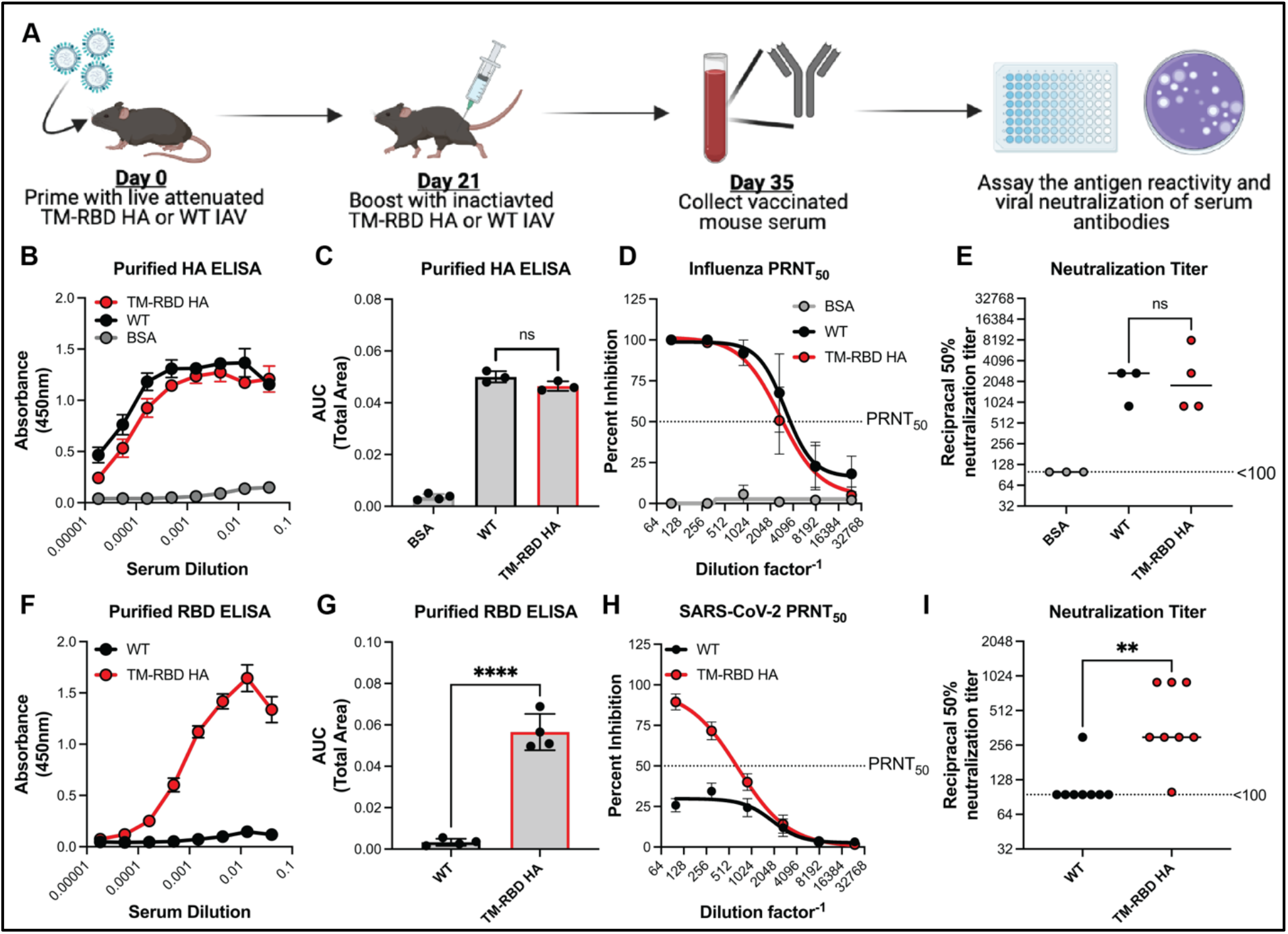
Vaccination of mice with the TM-RBD HA virus elicits strong neutralizing antibody responses to both IAV and SARS-CoV-2. (A) Experimental design displaying vaccination strategy, sample collection, and downstream assays. (B) ELISAs using serum from vaccinated mice against purified soluble HA protein. (C) Area under the curve analysis for (B). (D) IAV plaque reduction neutralization tests (PRNT) with serum from vaccinated mice. (E) Reciprocal 50% neutralization titer using IAV plaque neutralization data. (F) ELISAs using serum from vaccinated mice against purified soluble SARS-CoV-2 RBD protein. (G) Area under the curve analysis for (B). (H) SARS-CoV-2 plaque reduction neutralization tests (PRNT) with serum from vaccinated mice. (I) Reciprocal 50% neutralization titer using SARS-CoV-2 plaque neutralization data. Statistical analyses were performed using unpaired t-tests. For all panels, P-values denoted with asterisks correspond to the following values: ** < 0.01, and **** < 0.0001.

We observed high serum IgG reactivity against the IAV HA protein in both experimental vaccine groups but not the BSA control (**Figure 3B**). Further, there was no significant differences in the reactivity against HA between the two viral vaccine groups, despite there presumably being slightly less HA in the IAV/SARS-CoV-2 combination vaccine preparation (**Figure 3C**). For both experimental vaccine groups, the high antibody binding reactivity was correlated with high antibody neutralizing activity against authentic, infectious H1N1 IAV (**Figure 3D**). The average 50% plaque reduction neutralization titer (PRNT50) was calculated to be 1:3236 for WT and 1:2767 for TM-RBD HA, however those values were not significantly different from each other (**Figure 3E**).

Mouse immune serum from the combination viral vaccine group also showed strong reactivity against the SARS-CoV-2 RBD, with no notable reactivity from the WT IAV vaccine groups (**Figure 3F,G**). Plaque reduction assays with SARS-CoV-2 showed that while sera from WT IAV vaccinated animals had no appreciable neutralizing ability, the sera from the IAV/SARS-CoV-2 combination vaccine had strong neutralizing activity with a PRNT50 of approximately 1:679 (**Figure 3H,I**). Thus, the combination viral IAV/SARS-CoV-2 vaccine is immunogenic and elicits a potentially protective humoral response against SARS-CoV-2. This additional antigenicity comes at no apparent cost for IAV directed immune responses, validating the platform concept of a combination viral particle-based vaccine

### The IAV/SARS-CoV-2 combination vaccine induces a protective immune response against lethal challenges with IAV or SARS-CoV-2

Finally, we wanted to determine if the neutralizing antibody responses would also be associated with protection from influenza or SARS-CoV-2 induced disease. We therefore repeated the vaccination schemes and subsequently challenged mice with lethal doses of either IAV or SARS-CoV-2 (**Figure 4A**). For the IAV challenge, we performed an intranasal infection of C57BL/6 mice with the vaccine matched H1N1 strain A/Puerto Rico/8/1934 (PR8). While the BSA control vaccinated animals rapidly lost weight and succumbed to infection, both the WT IAV and the recombinant IAV/SARS-CoV-2 vaccine group were fully protected from virus-induced weight loss or mortality (**Figure 4B,C**).

**Figure 4.**
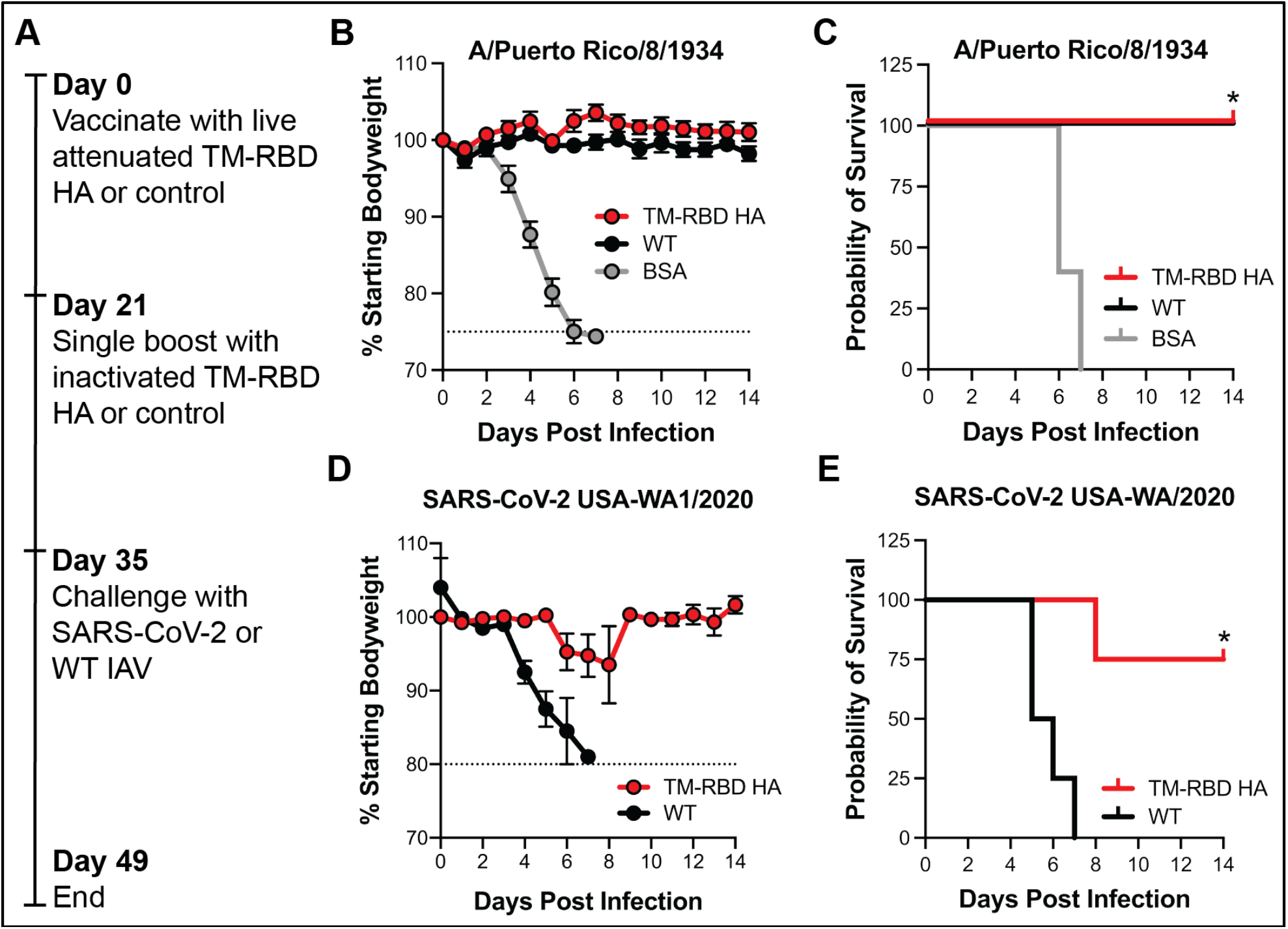
Mice vaccinated with TM-RBD HA virus are protected from lethal disease after challenge with IAV and SARS-CoV-2. (A) Diagram illustrating vaccination strategy and lethal challenges using IAV or SARS-CoV-2. (B) Bodyweight measurements of mice vaccinated with TM-RBD HA virus, WT IAV virus, or BSA and subsequently challenged with a lethal dose of A/Puerto Rico/8/1934 virus. (C) Survival of mice from (B). (D) Bodyweight measurements of mice vaccinated with TM-RBD HA virus or WT IAV virus and subsequently challenged with a lethal dose of USA-WA1/2020 SARS-CoV-2 virus. (E) Survival of mice from (D). Statistical analyses were performed using unpaired t-tests. For all panels, asterisks (*) indicate a P-value < 0.05.

In parallel, we also performed the same prime/boost regiment on transgenic Krt18-hACE2 mice on the C57BL/6 genetic background^25^. These mice express the human SARS-CoV-2 receptor and are thus infectible with WT SARS-CoV-2^26^. Two-weeks after the boost, a lethal dose of SARS-CoV-2^27^ was administered intranasally. While the control mice vaccinated with the WT IAV rapidly lost weight and succumbed to infection, the mice vaccinated with the combination IAV/SARS-CoV-2 vaccine were significantly more protected from viral disease (**Figure 4D,E**). Thus, at least in mouse models, vaccination with IAV-based SARS-CoV-2 combination vaccine can induce protective immunity, likely via the generation of neutralizing antibodies.

## Discussion

The development and production of novel vaccines is frequently labor intensive, costly, and difficult to scale upon high demand. The existing infrastructure for producing influenza vaccines is highly optimized and capable of delivering more than a billion doses per year^28^. In order to potentially leverage influenza vaccine infrastructure for the production of other vaccines, we have demonstrated that a lab adapted vaccine backbone (A/Puerto Rico/8/1934) can be genetically manipulated to express and package the receptor binding domain from the spike protein of SARS-CoV-2. Vaccination of mice with this recombinant virus elicited a high level of immunity and protected against lethal challenge with both IAV and SARS-CoV-2. These data together show that genetic engineering of influenza viruses may be a feasible platform approach to generate combination seasonal vaccines produced in a manner analogous to standard influenza vaccines.

Our data are in general agreement with previous work that has shown recombinant influenza viruses can be used as a vector to express other antigens. From expressing additional influenza proteins^22, 29, 30^ to expression of antigens and/or epitopes for pathogens as divergent as *Mycobacterium tuberculosis*^31^ and *Chlamydia trachomatis*^32^, the IAV can be modified to help elicit non-influenza directed responses from the immune system. In further support of this idea, another recombinant influenza-based strategy that does not package SARS-CoV-2 proteins on viral particles, but expresses them on the surface of cells during infection, can also induce SARS-CoV-2 antibodies^33^. Thus, data from multiple groups in addition to ours show that recombinant influenza viruses, delivered either as live attenuated or inactivated preparations, are a viable strategy for combination vaccine development against SARS-CoV-2 and other pathogens.

Despite the data presented in this study, there remain a number of questions that need to be answered. Perhaps most obvious question is if the S protein RBD is the most appropriate antigen relative to either the full S protein or other coronaviral proteins. While clearly vaccine responses against the RBD are sufficient to mediate some protection, it will be important to determine if boosting only RBD directed responses is ultimately an efficacious strategy in people. Further, our genetic manipulation of the virus was associated with a reduction in viral yield. Additional studies will be required to understand how decreased yield would affect the eventual vaccine cost or timelines for manufacturing. Finally, we have only utilized one genetic background and corresponding set of viral glycoproteins for these studies. While PR8 is a standard background for vaccine development, it will also be important to understand if other H1 (or indeed, H3 or influenza B virus) HA proteins are amenable to our manipulations to segment length and composition such that a membrane anchored RBD domain can be functionally encoded.

In conclusion, the influenza-based, multi-valent vaccines may represent a generalizable approach to reduce the time and manufacturing requirements for development of novel vaccines. Since the current influenza vaccine is composed of three or four distinct strains^34^, this approach could be even more highly multiplexed than eliciting responses against two pathogens. While there remain questions to be answered and technical challenges to overcome, “reprogramming” influenza viruses may be an attractive approach to produce and package antigens that are difficult to purify or are poorly immunogenic on their own. Continued work on this and other generalizable vaccine platforms will not only help with the current response to the COVID-19 pandemic but will help poise us for rapid response during future epidemic/pandemic outbreaks.

## Acknowledgements

The authors would like to thank Bart Haynes, Kevin Saunders, and Dapeng Li for their kind gifts of the DH1041 and DH1044 antibodies. This research has been funded in whole or part with federal funds under a contract from the National Institute of Allergy and Infectious Diseases, National Institutes of Health, Contract Number 75N93019C00050. The views, opinions and/or findings expressed are those of the authors and should not be interpreted as representing the official views or policies of the U.S. Government. The following reagent was deposited by the Centers for Disease Control and Prevention and obtained through BEI Resources, NIAID, NIH: SARS-Related Coronavirus 2, Isolate USA-WA1/2020, NR-52281. Biosafety level three SARS-CoV-2 studies were performed in the Duke Regional Biocontainment Laboratory, which received partial support for construction from the National Institutes of Health, National Institute of Allergy and Infectious Diseases (UC6-AI058607). Duke University may file for intellectual property protection of the viral genetic manipulation approaches in this manuscript.

## Methods

### Cell lines and viruses

Human embryonic kidney cells (HEK293T, ATCC) were grown in Dulbecco’s Modified Eagle Medium supplemented with 5% fetal bovine serum, GlutaMAX (Gibco cat. no. 35050079), and penicillin/streptomycin. Madin-Darby canine kidney cells (MDCK, ATCC) were grown in minimum essential medium (MEM) supplemented with 5% fetal bovine serum (FBS), GlutaMAX, HEPES, NaHCO_3_, and penicillin/streptomycin. All cells were grown at 37 °C under 5% CO_2_. African Green monkey kidney cells (Vero E6, ATCC) were grown in MEM+ Earl’s Salts + L-Glutamine (Gibco 11095-080). This media was supplemented with penicillin/streptomycin, 10% FBS, 1mM Na Pyruvate, and 1x MEM NEAA (Gibco 11140-050). A/Puerto Rico/8/1934 (PR8) virus was used for recombinant virus generation as well as vaccination/animal challenge experiments. SARS-Related Coronavirus 2, Isolate USA-WA1/2020, NR-52281 was used for SARS-CoV-2 infections and is from BEI Resources.

### Cloning and rescue of A/Puerto Rico/8/1934 encoding SARS-CoV-2 RBD

Influenza segment cloning was accomplished as previously described^22^. First, the receptor binding domain (RBD), which includes amino acids 319-541 of the spike protein open reading frame, from the SARS-CoV2 Wuhan-Hu-1 isolate (Accession: MN908947.3) was codon optimized for expression in influenza A viruses and synthesized by IDT. Importantly, we also encoded the neuraminidase (NA) transmembrane domain (amino acids 1-40 of the NA open reading frame) 5’ to the RBD to allow it to be incorporated into the viral particle, and a FLAG tag 3’ to the RBD to aid in detection as needed. Once this was done, the codon optimized RBD was PCR amplified and cloned into the bicistronic pDZ rescue plasmid system for A/Puerto Rico/8/1934 using the NEBuider HiFi DNA assembly kit (NEB). Specifically, the SARS-CoV-2 RBD-TM construct was cloned into the previously reported mNeon-HA construct, wherein the RBD-TM sequence replaced the mNeon reporter allowing expression of the transgene 5’ to the HA ^22^. Successful cloning was then confirmed by Sanger sequencing. Viral rescue was then performed by transfecting the modified RBD-TM HA plasmid alongside the 7 plasmids encoding the other PR8 segments into 293Ts using the Mirus TransIT-LT1 reagent. Rescued virus was then amplified via inoculation of virus into 10-day-old embryonated chicken eggs (Charles River) at 37 °C for three days.

### Virus propagation and growth kinetics

Influenza virus titering was performed as previously reported^22^. Briefly, approximately 500 PFU of each plaque-purified stock was injected in 10-day old embryonated hen eggs purchased from Charles River Laboratories, Inc. and incubated for 72 hours at 37 °C. The allantoic fluid was then harvested from and titer determined via plaque assay on MDCK cells. This was accomplished by serially diluting allantoic fluid and then incubating MDCKs with 500 μL of the diluted sample for 1 hour at 37 °C. After incubation, the viral suspension was aspirated, agar overlay was applied, and cells were incubated at 37 °C for 48-72 hours depending on plaque size. Plaque assays were then fixed by adding 2mL of 4% paraformaldehyde solution and incubating overnight at room temperature in a fume hood. The next day paraformaldehyde was aspirated, and cells were washed prior to performing antigen staining to detect A/Puerto Rico/8/1934 HA protein as described above. For viral growth curves and endpoint titer assays, eggs were injected with 10,000 pfu/egg and eggs were collected for plaque assay at 24, 48, and 72 hours post infection. Hemagglutination assays were performed by serially diluting virus 1: 2 in PBS in a 96 well plate then adding chicken blood to each well. Plates were incubated 1 hour then each well was scored as positive or negative. SARS-CoV-2 stocks were grown on Vero E6 cells in virus infection media (MEM+Earl’s Salts, penicillin/streptomycin, 2% FBS, 1mM Na Pyruvate, 1x MEM NEAA) for 72 hours. Stocks were frozen at −80 °C and were titered by serially diluting virus in virus infection media then infecting a confluent monolayer of Vero E6 cells growing in 6-well, poly-L-lysine treated, plates for 1 hour. Inoculum was then removed and an agarose overlay was added. Cells were incubated at 37 °C and 5% CO_2_ for 72 hrs then were stained with 0.05% neutral red in PBS for 3 hours.

### Viral purification

Purification of influenza viral particles was performed prior to use in vaccination and ELISA experiments. First, viral stocks were grown in 10-day old embryonated hen eggs as described above. Then allantoic fluid was collected and dialyzed overnight using the Spectra-Por Float-a-lyzer G2 10 mL, 100 kDA MWCO tubes according to manufacturer’s instructions (Millipore Sigma cat. no. Z727253-12EA). After samples were dialyzed to remove larger impurities, the allantoic fluid was collected and virus samples concentrated by ultracentrifugation using a 30% sucrose cushion for 1 hour at 25,700 RPM using the Sorvall TH-641 swinging bucket rotor. Virus samples were then resuspended in PBS and pooled prior to being fixed in 0.02% formalin for 30 minutes at room temperature. Samples were then once again dialyzed overnight to remove formalin using Slide-A-Lyzer casettes (ThermoScientific cat. no. PI66370) before being stored at 4 °C until use.

### RT-PCR and serial passage experiments

Viral RNA was extracted using TRIzol (Invitrogen) followed by cholorform/ethanol precipitation or using Qiagen viral RNA miniprep kits. RT-PCR reactions were performed using SuperScript™ III One-Step RT-PCR System kits (Thermo cat. no. 12574026) according to manufacturer’s guidelines. Primers used were, HA 5’ Forward: 5’-GTAGATGCAGCAAAAGCAGGGGAAAATA-3’, HA 3’ Reverse: 5’-CCATCCTCAATTTGGCAC-3’. RT-PCRs using RNA generated from miniprep kits varied in template concentration as the presence of carrier RNA inhibits nucleic acid quantification via photometric means. RT-PCR products were analyzed on 0.8-1% agarose gels run at 50V. For serial passage experiments, an 80% confluent monolayer of MDCK cells was infected with a MOI of 0.01 for 48 hrs. After 48 hrs, cell supernatants were collected and centrifuged for 5 minutes at 500 x g. Supernatant was removed and 100 μl was added to 1ml of Trizol and frozen at −80 °C. The titer of the remaining virus was then estimated by Hemagglutination assay and a new monolayer of cells was subsequently infected. This protocol was repeated 10 times.

### Microscopy

MDCK cells were seeded in 6-well plates coated with poly-L-lysine and allowed to grow at 37 °C for 24 hours prior to infection. Cells were washed with PBS and infected with either PR8 or PR8-TM-RBD-HA virus at an MOI of 1.0 in infection media (1 mM KH_2_PO_4_, 155 mM NaCl, Na_2_HPO_4_, 83.5 mM CaCl_2_, 105 mM MgCl_2_, 10 U/mL penicillin/streptomycin, 0.4% BSA) for 1 hour at 37 °C. Virus was removed from cells and replaced with post-infection media (Gibco OptiMem supplemented with 0.01% FBS, 10 U/mL penicillin/streptomycin, 0.4% BSA, and 1 μg/mL TCPK-trypsin). Cells were incubated for 24 hours and fixed with methanol-free 4% paraformaldehyde. DNA was visualized using Hoechst 33342 (Thermo) at 5 μg/mL in PBS for 15 minutes. HA and SARS-CoV-2 were detected using PY102 and SARS-CoV-2 Spike Protein (RBD) Polyclonal Antibody (Thermo cat. no. PA5-114451) at a 1:250 dilution for approximately 3 hours. Primary antibodies were visualized using AlexFluor594-conjugated anti-mouse/anti-rabbit secondary antibodies (Thermo cat. no. A-11032/R37117) at a dilution of 1:500 for 1 hour at room temperature.

### Western blots

Protein extracts were quantified and normalized via Bradford assay. SDS-PAGE was performed using 4-20% polyacrylamide gels (BioRad) electrophoresed at 120V for 60 minutes. Proteins were transferred to 0.45 μm nitrocellulose membranes at 90V for 60 minutes at 4 °C and blocked using PBST + 5% milk for a minimum for 1 hour at room temperature. For cellular lysates, 20 μg of total protein was loaded per sample. To normalize viral protein extracts, 0.5 μg of PR8 and PR8-TM-RBD HA were initially loaded and analyzed via western blot. Viral protein extracts were probed for M1 and normalized via densitometry (ImageJ). After normalization to M1, 0.5 μg PR8 and 1.32 μg PR8-TM-RBD HA were loaded for subsequent western blot analyses. The following antibodies were used for protein detection; PY102 (HA, 1 μg/mL), 4A5 (NA, 0.45 μg/mL), anti-matrix protein [E10] (M1 and M2, 1:1,000, Kerafast cat. no. EMS009), and anti-SARS-CoV-2 spike protein (RBD) polyclonal antibody (RBD, 1:1,000, Invitrogen cat. no. PA5-114451). All primary antibodies were diluted in PBST + 5% milk and applied to membranes for ≥16 hours at 4 °C. Anti-mouse (1:20,000, Thermo cat. no. A16072) and anti-rabbit (1:10,000, Thermo cat. no. A16104) secondary antibodies were diluted in PBST + 5% milk and applied to membranes for 60 minutes at room temperature. Membranes were developed using Clarity ECL or Clarity ECL MAX.

### ELISAs

Proteins/viruses were immobilized to 96-well plates using carbonate coating buffer (30 mM Na2CO3, 70 mM NaHCO3, pH 9.5) for ≥16 hours at 4 °C. For protein samples, 50-100 μL of sample at 10 μg/mL was added to wells. For viral samples, 1 x 10^6^ PFUs were added to wells in a volume of 50-100 μL. All samples were diluted using PBS + 3% BSA. After coating, wells were washed with PBS and blocked with PBS + 3% BSA for at least 1 hour at room temperature. Primary antibodies were diluted using PBS + 3% BSA and incubated with immobilized proteins/viruses for ≥1 hour at room temperature (anti-RBD ProSci cat. no. 9087, anti-HA PY102 a kind gift from Dr. Tom Moran (Icahn School of Medicine at Mount Sinai), DH1041 and DH1044 kind gift from Drs. Bart Haynes, Kevin Saunders, and Dapeng Li (Duke)). Anti-human (1:10,000 Thermo cat. no. A18805), anti-mouse (1:5,000, Thermo cat. no. A16072), and anti-rabbit (1:5,000, Thermo cat. no. A16104) secondary antibodies were diluted in PBS + 3% BSA and incubated with immobilized proteins/virus for 1 hour at room temperature. ELISAs were developed using 1-Step Ultra TMB-ELISA substrate (Thermo cat. no. 34029) and quenched with 2M H2SO4.

### Animal infections/Live-attenuated vaccination

Animal infections were performed using age-matched C57BL/6J (Jackson Labs 000664) or B6.Cg-Tg(K18-ACE2)2Prlmn/J mice (Jackson Labs 034860) in accordance with Duke University IACUC approved protocol numbers A189-18-08 and A081-20-04. For influenza infections, mice were anesthetized using 100 μL of a 14.2 mg/ml ketamine-xylazine mixture via intraperitoneal injection. After administration of anesthesia, mice were marked and baseline weights were measured. Mice were then given a 40 μL intranasal inoculum of virus, at the 10 pfu WT PR8 or 250 pfu PR8 TM-RBD HA, diluted in pharmaceutical-grade PBS. After infection, mice were weighed daily and euthanized once their body weight reached 75% of the baseline weight for IAV. This cutoff was determined respective to the weight prior to infection. Euthanasia was performed via a primary method of CO_2_ asphyxiation and bilateral thoracotomy as a secondary confirmation. These methods were used during both challenge experiments and administration of the live virus vaccine prime. For SARS-CoV-2 challenge experiments, prior to infection, mice were injected subcutaneously with IPT-300 transponders capable of reading body temperature and animal ID (BMDS IPT-300) and baseline weights and temps were measured. On the day of infection, mice were anesthetized using isoflurane then given a 50 μL intranasal inoculum of virus, 3×10^4^ pfu, diluted in pharmaceutical-grade PBS. After infection, mice were monitored daily for weight, temperature and clinical signs. Mice were euthanized via CO_2_ asphyxiation and bilateral thoracotomy as a secondary confirmation when their body weight reached 80% of the baseline weight measured prior to infection or reached a clinical score of 4 in accordance with the approved protocol above A081-20-04.

### Vaccination of mice

Vaccine doses for the boost were prepared using the purified inactivated virus described above. Virus samples were then diluted 1:1 with the adjuvant Addavax (Invivogen cat. no. vac-adx-10) to a final concentration of 100 μg/mL. After preparation of doses, mice were anesthetized as described above and then administered a 100 μL injection intramuscularly into the left leg. Mice were then monitored the next day for side effects and then housed for the indicated period of time before collection or viral challenge. If serum was collected, mice were administered a lethal dose of the ketamine-xylazine (200 μL) and then blood was harvested and serum collected using Sarstedt Z-Gel tubes according to manufacturer’s instruction (Sarstedt cat. no. 41.1378.005). Serum was then stored at −20 °C until use.

### Plaque reduction neutralization assays

MDCK and Vero cells were used for influenza and SARS-CoV-2 plaque reduction assays, respectively. A master mix of virus was diluted to the indicated concentration (~40-80 PFU/mL) and aliquoted prior to being mixed with antibody dilutions. Following a 45-minute incubation at room temperature with antibody, the media was aspirated from cells and 500 μL of the virus-antibody mixture was added to each well of cells. For each experiment a no antibody control was included to accurately record how much virus had been used to infect cells. Cells were incubated with the virus-antibody mixture at 37 °C for 1 hour, rocking the samples every 15 minutes to ensure cells were completely covered by the solution. After this period, the solution was aspirated, and an agar overlay was applied. For influenza plaque reduction assays, staining and plaque counting was performed as described above in the titering section. SARS-CoV-2 plaque reduction assays were evaluated by first staining plaques with .05% Neutral Red solution for 3 hours at 37 °C (Sigma Aldrich cat. no. N2889-100mL). Neutral red was then aspirated from the wells and plaques were counted. The percent reduction in plaques was calculated in reference to a no sera control. The PRNT50 value was calculated by using a non-linear regression model, [Agonist] vs. response—Find ECanything with f constrained to 50. The reciprocal 50% neutralization titer was calculated by averaging the greatest dilution of mouse sera that had a >50% reduction in plaques compared to a no sera control for each sera sample in a vaccination group.

## References

1. Harding AT, Heaton NS. Efforts to Improve the Seasonal Influenza Vaccine. Vaccines (Basel) 6, (2018).

2. Trombetta CM, Gianchecchi E, Montomoli E. Influenza vaccines: Evaluation of the safety profile. Hum Vaccin Immunother 14, 657–670 (2018).

3. Savidan E, Chevat C, Marsh G. Economic evidence of influenza vaccination in children. Health Policy 86, 142–152 (2008).

4. Sullivan SG, Price OH, Regan AK. Burden, effectiveness and safety of influenza vaccines in elderly, paediatric and pregnant populations. Ther Adv Vaccines Immunother 7, 2515135519826481 (2019).

5. Nogales A, Martinez-Sobrido L. Reverse Genetics Approaches for the Development of Influenza Vaccines. Int J Mol Sci 18, (2016).

6. Fiege JK, Langlois RA. Investigating influenza A virus infection: tools to track infection and limit tropism. J Virol 89, 6167–6170 (2015).

7. Dumm RE, Heaton NS. The Development and Use of Reporter Influenza B Viruses. Viruses-Basel 11, (2019).

8. Jenkins MR, Webby R, Doherty PC, Turner SJ. Addition of a prominent epitope affects influenza a virus-specific CD8(+) T cell immunodominance hierarchies when antigen is limiting. Journal of Immunology 177, 2917–2925 (2006).

9. Hu B, Guo H, Zhou P, Shi ZL. Characteristics of SARS-CoV-2 and COVID-19. Nat Rev Microbiol 19, 141–154 (2021).

10. Dong Y, Dai T, Wei Y, Zhang L, Zheng M, Zhou F. A systematic review of SARS-CoV-2 vaccine candidates. Signal Transduct Target Ther 5, 237 (2020).

11. Sadoff J, et al. Interim Results of a Phase 1-2a Trial of Ad26.COV2.S Covid-19 Vaccine. N Engl J Med, (2021).

12. Baden LR, et al. Efficacy and Safety of the mRNA-1273 SARS-CoV-2 Vaccine. N Engl J Med 384, 403–416 (2021).

13. Polack FP, et al. Safety and Efficacy of the BNT162b2 mRNA Covid-19 Vaccine. N Engl J Med 383, 2603–2615 (2020).

14. Rubin R. COVID-19 Vaccines vs Variants-Determining How Much Immunity Is Enough. Jama-J Am Med Assoc, (2021).

15. Honda-Okubo Y, Barnard D, Ong CH, Peng BH, Tseng CT, Petrovsky N. Severe acute respiratory syndrome-associated coronavirus vaccines formulated with delta inulin adjuvants provide enhanced protection while ameliorating lung eosinophilic immunopathology. J Virol 89, 2995–3007 (2015).

16. Poland GA, Ovsyannikova IG, Kennedy RB. SARS-CoV-2 immunity: review and applications to phase 3 vaccine candidates. Lancet 396, 1595–1606 (2020).

17. Sariol A, Perlman S. Lessons for COVID-19 Immunity from Other Coronavirus Infections. Immunity 53, 248–263 (2020).

18. Huang AT, et al. A systematic review of antibody mediated immunity to coronaviruses: kinetics, correlates of protection, and association with severity. Nat Commun 11, 4704 (2020).

19. Lan J, et al. Structure of the SARS-CoV-2 spike receptor-binding domain bound to the ACE2 receptor. Nature 581, 215-+ (2020).

20. Tai WB, et al. Characterization of the receptor-binding domain (RBD) of 2019 novel coronavirus: implication for development of RBD protein as a viral attachment inhibitor and vaccine. Cell Mol Immunol 17, 613–620 (2020).

21. Soema PC, Kompier R, Amorij JP, Kersten GF. Current and next generation influenza vaccines: Formulation and production strategies. Eur J Pharm Biopharm 94, 251–263 (2015).

22. Harding AT, Heaton BE, Dumm RE, Heaton NS. Rationally Designed Influenza Virus Vaccines That Are Antigenically Stable during Growth in Eggs. Mbio 8, (2017).

23. Selzer L, Su ZM, Pintilie GD, Chiu W, Kirkegaard K. Full-length three-dimensional structure of the influenza A virus M1 protein and its organization into a matrix layer. Plos Biol 18, (2020).

24. Li D, et al. The functions of SARS-CoV-2 neutralizing and infection-enhancing antibodies in vitro and in mice and nonhuman primates. bioRxiv, (2021).

25. McCray PB, Jr., et al. Lethal infection of K18-hACE2 mice infected with severe acute respiratory syndrome coronavirus. J Virol 81, 813–821 (2007).

26. Winkler ES, et al. SARS-CoV-2 infection of human ACE2-transgenic mice causes severe lung inflammation and impaired function (vol 51, pg 831, 2020). Nat Immunol 21, 1470–1470 (2020).

27. Winkler ES, et al. SARS-CoV-2 infection of human ACE2-transgenic mice causes severe lung inflammation and impaired function. Nat Immunol 21, 1327–1335 (2020).

28. Sparrow E, et al. Global production capacity of seasonal and pandemic influenza vaccines in 2019. Vaccine 39, 512–520 (2021).

29. Pena L, et al. Influenza Viruses with Rearranged Genomes as Live-Attenuated Vaccines. Journal of Virology 87, 5118–5127 (2013).

30. Gao Q, Lowen AC, Wang TT, Palese P. A nine-segment influenza a virus carrying subtype H1 and H3 hemagglutinins. J Virol 84, 8062–8071 (2010).

31. Sereinig S, et al. Influenza virus NS vectors expressing the mycobacterium tuberculosis ESAT-6 protein induce CD4+ Th1 immune response and protect animals against tuberculosis challenge. Clin Vaccine Immunol 13, 898–904 (2006).

32. He Q, et al. Live-attenuated influenza viruses as delivery vectors for Chlamydia vaccines. Immunology 122, 28–37 (2007).

33. Loes AN, Gentles LE, Greaney AJ, Crawford KHD, Bloom JD. Attenuated Influenza Virions Expressing the SARS-CoV-2 Receptor-Binding Domain Induce Neutralizing Antibodies in Mice. Viruses 12, (2020).

34. Wei CJ, Crank MC, Shiver J, Graham BS, Mascola JR, Nabel GJ. Next-generation influenza vaccines: opportunities and challenges. Nat Rev Drug Discov 19, 239–252 (2020).

